# A Propidium Iodide-based *In Vitro* Screen of the “Bug Box” Against *Babesia duncani* Reveals Potent Inhibitors

**DOI:** 10.1101/2025.01.14.633027

**Authors:** Emmett A. Dews, José E. Teixeira, Christopher D. Huston, Marvin J. Meyers, Peter R. Hyson

## Abstract

Incidence and endemic range of human babesiosis are expanding. Standard therapy for human babesiosis consists of antimicrobials developed for other indications. While these treatments are adequate in immunocompetent hosts, infections in the immunocompromised can be severe, relapsing, and drug-resistant despite use of multi-drug regimens. Existing drugs are ineffective because they cannot safely achieve and maintain adequate serum concentrations to inhibit Babesia. Discovery of improved agents against Babesia spp. is of growing importance and efficient techniques for high throughput compound screening can assist in this effort.

We developed a high throughput *in vitro* drug screening assay for *Babesia duncani* that is conducted in 384 well plates and makes use of the fluorescent DNA stain propidium iodide (PI) with relative fluorescence measured by a microplate reader. A Z’ factor of >0.81 was calculated which suggests an excellent ability to detect inhibitory compounds. A screen of the 41-compound library Structural Genomics Consortium Bug Box was conducted yielding five hits: trimethoprim, atovaquone, SDDC M7, diphenyleneiodonium chloride, and panobinostat. Panobinostat, a histone deacetylase complex (HDAC) inhibitor, was selected for further evaluation given that its target had not been previously explored in *B. duncani.* Follow up dose-response testing of structurally related compounds revealed multiple potential leads including nanatinostat and quisinostat, both of which were potent at the nanomolar level and showed favorable selectivity index in cytotoxicity studies. High throughput screening using PI and 384 well plates is an advance in drug discovery for babesiosis and HDAC inhibitors show promise as lead compounds worthy of further investigation.

## BACKGROUND

Babesiosis, the disease caused by the intracellular parasite *Babesia*, is an emerging infectious disease with rising incidence and expanding endemic range(1–3). *Babesia* are tickborne, malaria-like protozoa of the phylum Apicomplexa which infect red blood cells of vertebrates including humans. The spectrum of human disease is wide with many infections asymptomatic or causing a mild, self-limited, flulike illness but a few-particularly in patients at the extremes of age or with compromised immune systems-leading to critical illness and even death(4). Patients with asplenia are at particularly high risk of fulminant disease due to inability to clear the bloodstream of infected erythrocytes.

Standard of care antimicrobial treatment for Babesiosis consists of combination therapy using clindamycin/quinine or atovaquone/azithromycin, antimicrobials developed and dosed for other indications such as malaria(5). While this treatment is generally effective in immunocompetent patients, in immunocompromised hosts infections can be relapsing and even incurable despite use of multi-drug regimens and antimicrobial resistance can develop during therapy(6–8). Indeed, evidence from animal models and *in vitro* culture suggests that most of the standard antimicrobial agents used for this indication have poor efficacy with minimum inhibitory concentrations (MICs) orders of magnitude higher in *Babesia* than *Plasmodium*, treatment failure in SCID mice, and rapid development of antimicrobial resistance on single agent therapy(9–12). Identification of new therapies against *Babesia* is of increasing concern given the expansion of this disease and the large and growing population of persons with asplenia and other immunocompromising conditions(13).

High throughput screening can be an efficient way to identify compounds that have activity for a particular indication and new drug targets that can be leveraged with rational drug design. Such an assay needs to be rapid, reproducible, capable of testing large numbers of compounds in small quantity, and capable of accurately distinguishing hits from compounds that lack significant activity. An *in vitro* drug screening assay using miniaturized cultures of *Babesia* in 96-well plates and SYBR green fluorescent labeling to identify infected erythrocytes has been developed and validated with *B. bovis* and utilized widely for multiple *Babesia* species(9, 14–16). To our knowledge, however, validation of such an assay in 384 well plates and using the inexpensive reagent propidium iodide (PI) has not been described. Use of the 384 well plate for high throughput screening is beneficial because it increases the number of compounds that can be simultaneously screened, uses less compound, and is more cost effective. *Babesia duncani* is an ideal organism to use for drug discovery using high throughput screening as it may both be continuously cultured in vitro (unlike the prevalent human pathogen *B. microti*) and because it has a murine model that may be used for further characterization and optimization of lead compounds(17). Herein we describe optimization of a high throughput, PI-based drug screening assay for *B. duncani* and the deployment of this assay in screening a collection of 41 compounds with reported antimicrobial activity that was previously assembled by the Structural Genomics Consortium(18). We report identification and characterization of novel classes of potent anti-*Babesia* compounds.

## RESULTS

### Propidium-iodide based drug assay optimization

To find a reproducible and cost-effective method to linearly correlate parasitemia to relative fluorescent units (RFU) in a 384 well plate, several fluorescent dyes were assessed. In a simple test of infected v. uninfected RBCs with the fluorescent dyes held at identical concentrations, propidium iodide (PI) produced a significant difference by Welch’s t-test and had the highest signal to background (Supplemental). It appeared superior even to the commonly used fluorescent stain SYBR green which produced more fluorescence in uninfected than infected cells. Diluting parasitemia experiments were performed iteratively in order to optimize the PI staining protocol. Fig 1A illustrates a close correlation between fluorescence from PI staining and parasitemia as shown in two independent experiments with R^2^>0.95.

**FIG 1.**
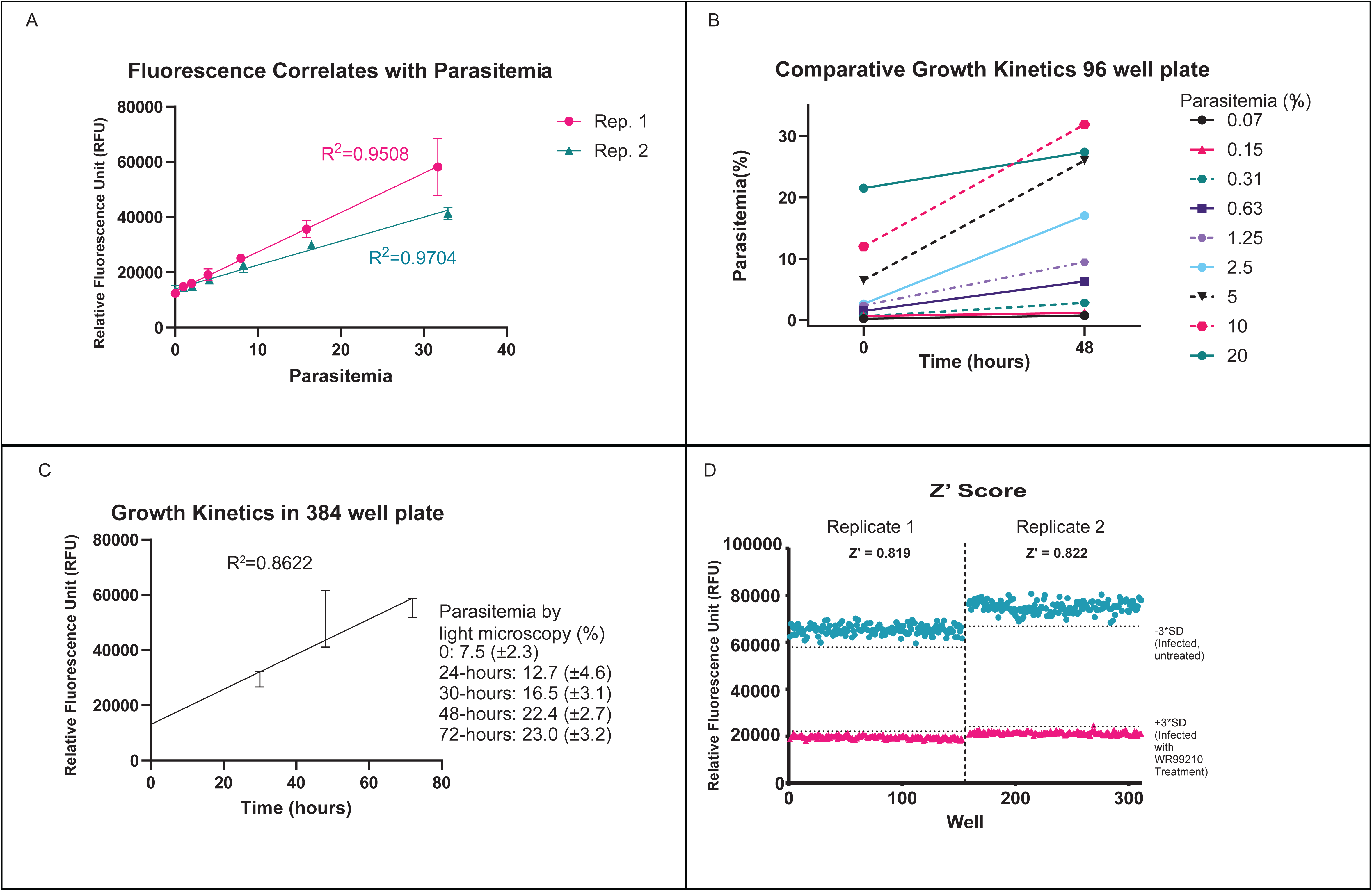
(**A)**Serial dilutions of high parasitemia *B. duncani* were made and seeded into a 384 well plate with three well technical replicates. Wells were stained with PI and then read using a microplate reader. Both repeats of this experiment are shown. **(B)** *B. duncani* infected red blood cells seeded into a 96 well plate at different starting parasitemias and incubated 48 hours. Parasites manually counted from giemsa-stained blood smears at time 0 and 48 hours. Each point represents parasitemia of a single well. **(C)** *B. duncani* infected red blood cells seeded into 384 well plates at parasitemia of 7.5% and then analyzed at 24, 30, 48, 72 hours with parasitemia measured using both PI staining (on graph) and blood smears and manual parasite counting (text in legend). time zero point included 22 wells, the 24-hour time point included 28 wells, and all others were 27 well technical replicates. **(D)** Half of a 384 well plate was seeded with *B. duncani* infected red blood cells and treated 48 hours with WR99210 while half was incubated with DMSO vehicle. Plate was then stained with PI and read with microplate reader. Z’ score calculated as described in methods. Both experiment replicates are displayed.

Growth kinetics experiments were also performed to assess the ability of *B. duncani* to propagate in miniaturized culture conditions. Fig 1B shows the 48 hour growth of *B. duncani* in a 96 well plate with multiple different starting parasitemias. Fig 1C shows the 72 hour growth of *B. duncani* in a 384 well plate as measured by PI staining and fluorescence with results of manually scoring blood smears indicated in text. It was established that in a culture of 5% hematocrit and with starting parasitemia of 7.5%, 48 hour parasitemia could be expected to be >20% in a 384 well plate. In a 96-well plate, parasitemia would more than triple in the course of 48 hours where starting parasitemia was 2.5-10%. Moreover, parasite growth could be accurately and reproducibly measured with PI staining and microplate fluorimetry.

### Assay validation for library screening

The Z or Z’-score, a statistical coefficient that describes an assay’s signal window, is commonly used to assess an assay’s suitability for compound library screening with Z-scores of 0.5 to 1 indicating an excellent assay(19). PI is expected to stain persistent DNA from killed or inhibited *Babesia.* We therefore determined the Z’-score by comparing fluorescence of untreated infected wells to infected wells treated with the dihydrofolate reductase inhibitor WR99210, a known inhibitor of *Babesia* growth(20). As shown in FIG1D, this experiment yielded a Z’ score of >0.81, consistent with an excellent ability to detect an effective inhibitory agent(19).

### Pilot Testing of Vehicle and Known Compounds

Having optimized staining protocols and culture conditions, dose response assays were conducted with DMSO vehicle, WR99210 (control agent with known inhibitory activity on *B. duncani*) and standard of care drugs atovaquone, azithromycin, clindamycin, quinine. Results are shown in Figure 2. DMSO, the vehicle in which experimental compounds are dissolved, lacked significant inhibitory activity at concentrations up to 0.5% by volume. WR99210 was demonstrated to have a 50% effective concentration (EC50) of 8.3 nM while atovaquone was also quite potent with EC50 of 518 nM. Azithromycin and quinine were only modestly effective with EC50s of 12.37 and 17.49 μM respectively. Clindamycin appeared not to inhibit at all.

**FIG 2.**
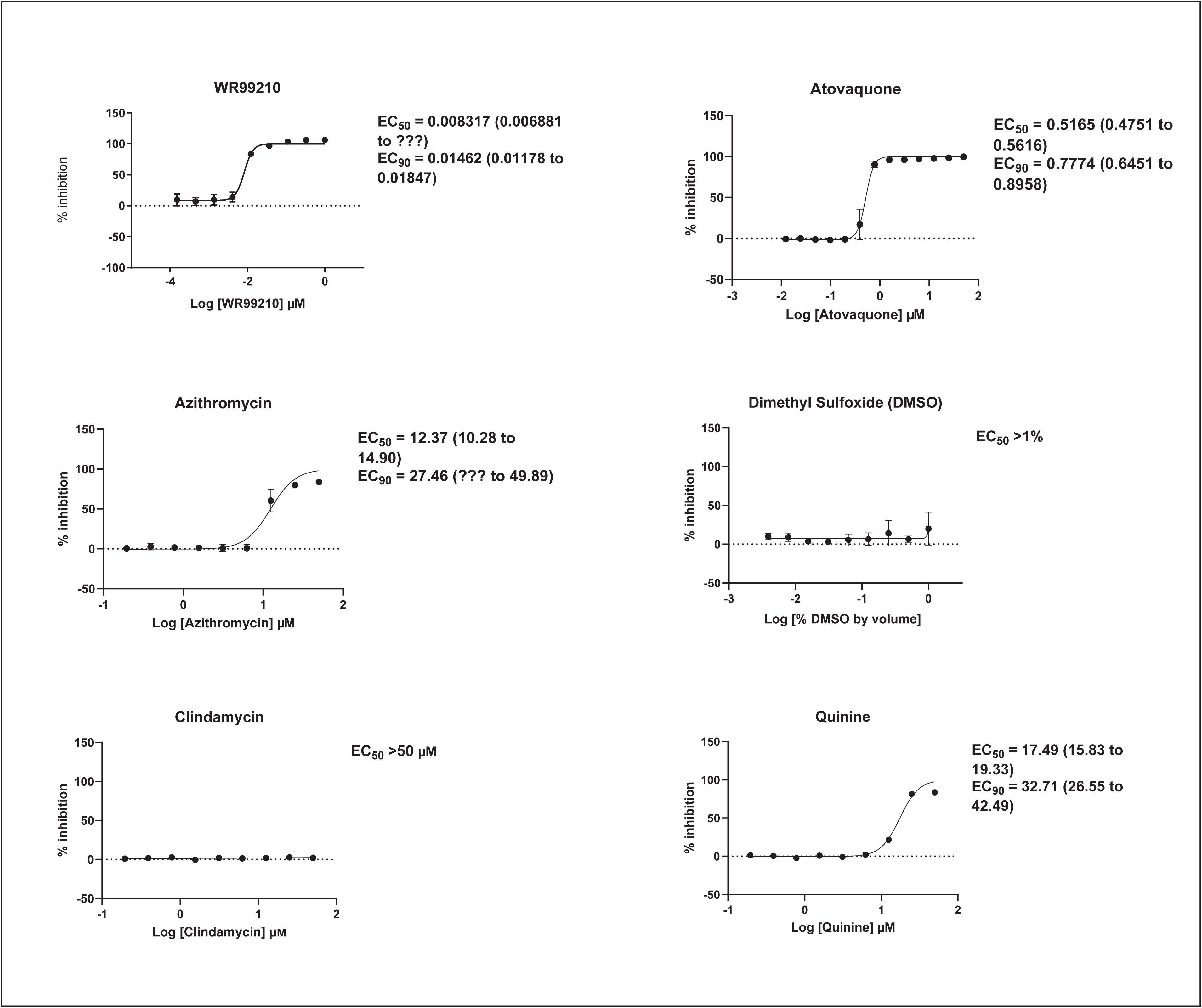
Dose response Curves for standard of care compounds and control compounds with *B. duncani* in vitro. Generated using nine point serial dilution of compound and treatment of parasite infected red bloods in 384 well plates with 48 hour incubation and propidium iodide staining. Four technical replicate wells per compound dilution and experiments repeated twice. WR99210 is a DHFR inhibitor commonly used in experiments with *Babesia*. DMSO is used as drug vehicle in our experiments.

### Pilot Screen & Hit Validation

The SGC previously assembled a small collection of known antimicrobial compounds (the “Bug Box”) that we used to test our assay in practice while identifying new anti-*Babesia* compounds(18). Compounds were screened at 5 μM and 1 μM, and compounds that inhibited greater than 80% were selected for re-supply and dose-response testing. Five compounds met the hit definition at 5 μM: panobinostat, trimethoprim, diphenylene iodonium chloride, atovaquone, and SDDC M7(Fig. 3A). Fig 3B shows results of dose-response testing with commercially resupplied compounds. Atovaquone, was not retested as it is known to be inhibitory toward *Babesia* and an EC50 had been generated previously. While trimethoprim proved only modestly effective with an EC50 of about 2.7 µM, the remaining 3 agents were quite potent-100% inhibitory in sufficient quantity. Of these, the histone deacetylase (HDAC) inhibitor panobinostat was most impressive with an EC50 of 8.4 nM.

**FIG 3.**
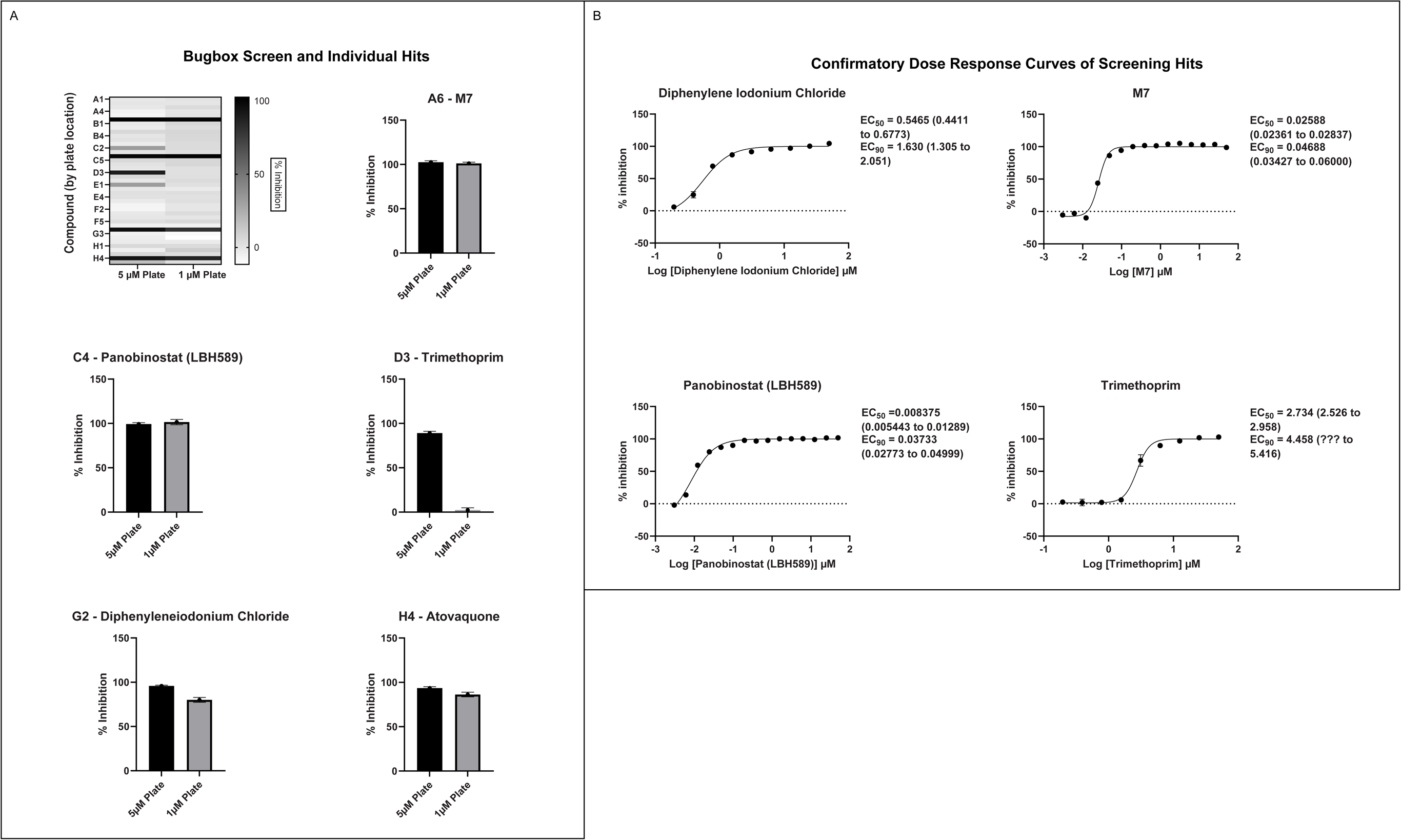
Bug Box Initial Screen and Hit validation **(A)**Screen of the Bug Box was run in two 384 well plates with one testing 1 µm dilutions and the other 5 µM compound dilutions with *B. duncani* infected red blood cells. Heatmap shows relative inhibitory activity by compound location in the plate with darker bars indicating greater inhibition and hits magnified. **(B)**Validation of hits with dose response curves using compound commercially obtained (with the exception of M7). Atovaquone excluded as it had been previously validated

### Histone Deacetylase Complex Inhibitor testing

In light of the high potency of panobinostat and the novelty of the putative target (HDAC) in *Babesia duncani,* we decided to test other HDAC inhibitors so as to begin elucidating SAR. A selection of commercially available HDAC inhibitors was chosen from among hydroxamates and benzamides which had been either approved for clinical use or had reached late stages of development(21). Dose response curves were generated with EC50s given in Table 1. All of the compounds tested proved to have inhibitory activity, most with EC50s in the nanomolar range. Quisinostat was the most potent with an EC50 of 4.5 nM. It was observed that unsaturated and aryl hydroxamic acids were more inhibitory than saturated vorinostat and that hydroxamic acids in general had more potent EC50s than did benzamides mocetinostat and entinostat.

**Table 1.**
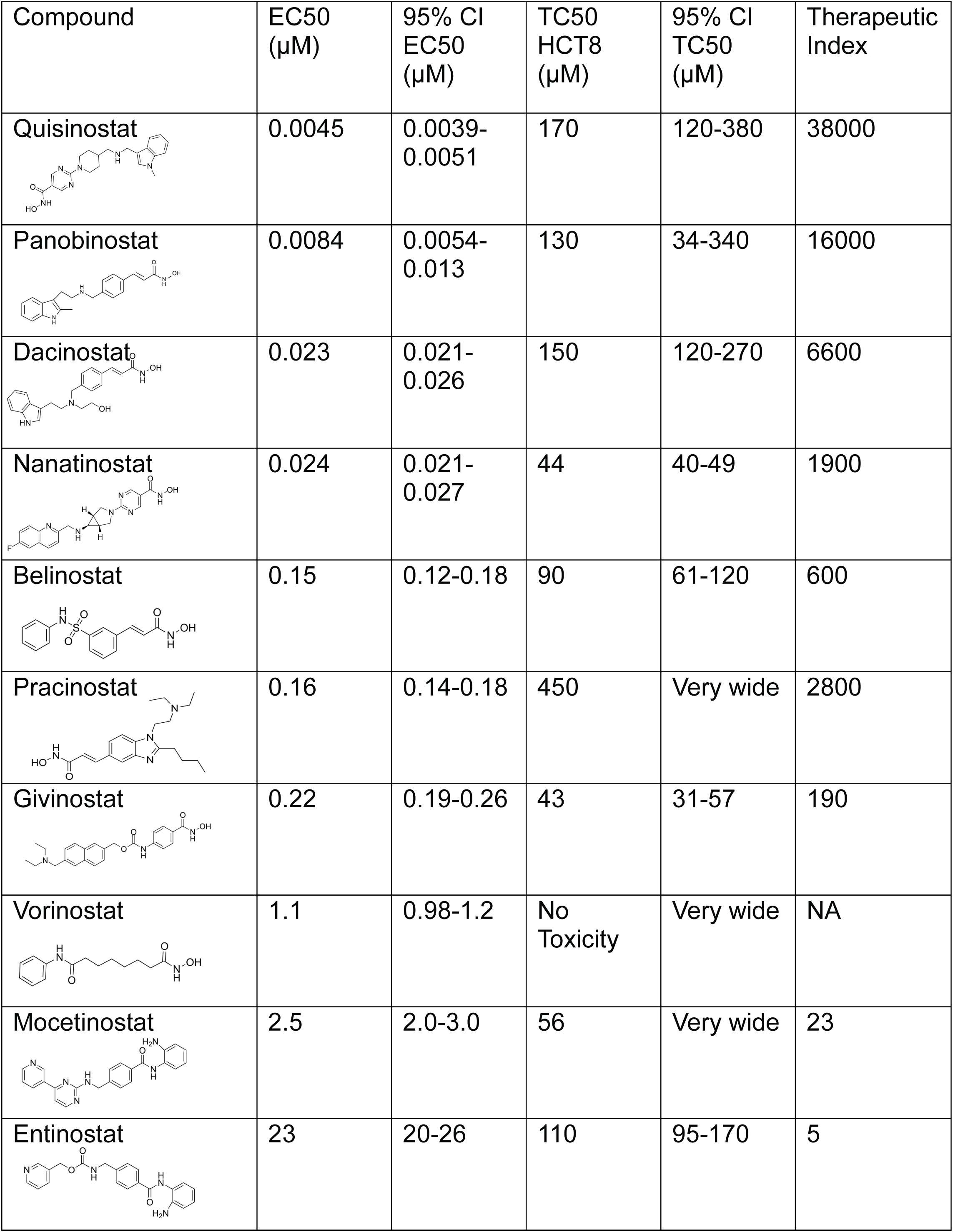
EC50s, cytotoxicity, structure activity relationships of select histone deacetylase inhibitors. Compounds are arranged by inhibitory potency against *B. duncani* with the most potent on top.

| Compound | EC50<br>(μM) | 95% CI<br>EC50<br>(μM) | TC50<br>HCT8<br>(μM) | 95% CI<br>TC50<br>(μM) | Therapeutic<br>Index |
| --- | --- | --- | --- | --- | --- |
| **Quisinostat** | 0.0045 | 0.0039-0.0051 | 170 | 120-380 | 38000 |
| **Panobinostat** | 0.0084 | 0.0054-0.013 | 130 | 34-340 | 16000 |
| **Dacinostat** | 0.023 | 0.021-0.026 | 150 | 120-270 | 6600 |
| **Nanatinostat** | 0.024 | 0.021-0.027 | 44 | 40-49 | 1900 |
| **Belinostat** | 0.15 | 0.12-0.18 | 90 | 61-120 | 600 |
| **Pracinostat** | 0.16 | 0.14-0.18 | 450 | Very wide | 2800 |
| **Givinostat** | 0.22 | 0.19-0.26 | 43 | 31-57 | 190 |
| **Vorinostat** | 1.1 | 0.98-1.2 | No<br>Toxicity | Very wide | NA |
| **Mocetinostat** | 2.5 | 2.0-3.0 | 56 | Very wide | 23 |
| **Entinostat** | 23 | 20-26 | 110 | 95-170 | 5 |

### Cytotoxicity Testing

Basic cytotoxicity testing of HDAC inhibitors was performed in human HCT8 cells as described above. Results are reported in Table 1. HDAC inhibitor compounds did show toxicity in human HCT8 cells but at concentrations significantly higher than what was required for inhibition of *B. duncani*. For example, while the compound most toxic to HCT8 cells appeared to be nanatinostat, which had an EC50 of 56.25 µM, by contrast its EC50 of 23 nM in *B. duncani* was 2400x lower.

## DISCUSSION

Effective treatment for human babesiosis is a public health concern of growing significance(22). *In vitro* testing of recommended therapies in *B. duncani* reported both here and elsewhere shows that of the four drugs used in clinical practice, only atovaquone would be considered significantly inhibitory with an EC50 of ∼500 nM. In the effort to discover new antimicrobials and druggable targets, a high throughput screening assay which is both cheaper and more efficient will be an invaluable tool. The assay described above uses a new parasite staining method and miniaturized culture conditions, and its Z’-score of 0.81 indicates its utility for library screening. Use of PI and the miniaturized culture format of 384-well plates will minimize the cost of reagents and the quantity of compounds required for screening and in vitro follow up, making this method an attractive alternative to those previously reported(9, 14–16). This assay’s utility in practice is demonstrated by its deployment in a screen of the SGC Bug Box compound collection with yield of both known and novel compounds inhibitory to *Babesia duncani*.

Diphenyleneiodonium chloride (DPI) is believed to inhibit NADP/NADPH oxidase and has been identified as an inhibitor of *M. tuberculosis, S. aureus*, and multiple species of non-Tuberculous mycobacteria(23, 24). One study synthesized two series of DPI analogues and showed good activity against *P. falciparum*, however significant concerns around cytotoxicity were noted(25). There are currently no published human trials with DPI. While this certainly may represent a potential chemical structure and/or target for further pursuit, the apparent lack of further progress in development of this compound for clinical use suggests that there are significant obstacles to its study.

Trimethoprim and SDDC-M7 both are thought to target parasite dihydrofolate reductase (DHFR), a critical enzyme for tetrahydrofolate synthesis required for synthesis of purines and some amino acids. Control compound WR99210 also targets DHFR(26, 27), and this drug target has been extensively studied and described in *Babesia* and other parasites(20). Unfortunately SDDC-M7 is not commercially available and the researchers who originally synthesized it could not be contacted, thus further study of this hit compound by our group was not possible.

The molecular target of panobinostat is believed to be a family of epigenetic modification enzymes called histone deacetylase complexes. Panobinostat has been approved as an oral treatment for multiple myeloma, and other HDAC inhibitors have been used for numerous forms of both solid organ and hematologic malignancy(28). There are 18 mammalian HDACs which are divided into two families: the histone deacetylase family and the sirtuin protein family(29). Panobinostat is considered a nonselective inhibitor within the histone deacetylase family. While its known activity in human cells (and role as a chemotherapy agent) could be considered a red flag, prior study has shown that panobinostat’s cytotoxicity is fairly selective for malignant cells, possibly because HDACs are relatively more expressed in this cell population(30). While panobinostat LD90s have been reported as less than 500 nM for solid tumor cells and less than 100 nM for liquid tumor cell lines, testing in normal cell lines (human mammary and renal epithelial cells among others) has shown LD90s >5μM. Similarly, while malignant cells treated with panobinostat had a 10-11 fold increase in apoptotic signaling, normal cells showed no increase in caspase expression compared with control(31). Cardiac toxicity has been an issue in this drug class, however medicinal chemistry optimization approaches have successfully yielded compound derivatives that mitigate this problem(21).

HDAC inhibitors have also been explored for therapeutic applications against intracellular parasites including other Apicomplexa such as *P. falciparum* and *T. gondii* with some displaying good selectivity for parasite over host cells and even activity against drug resistant *P. falciparum* (*32*). While apicidin has been shown to have inhibitory activity on *B. bovis*(*33*) and a screen of epigenetic modifying compounds showed inhibitory activity of HDAC inhibitors in *Babesia divergens*, it does not appear that such compounds have previously been tested in *Babesia duncani*(*34*).

From a chemistry perspective, most of the HDAC inhibitors evaluated, including compounds panobinostat, vorinostat, belinostat, dacinostat, are hydroxamic acids while entinostat and mocetinostat are benzamides(35). Our early-stage SAR indicates that the hydroxamic acids have greater inhibitory activity against *B. duncani* than do the benzamides. The greater anti-*Babesia* potency of the hydroxyamic acids vs. benzamides may related to differences in target selectivity, since the benzamides entinostat and mocetinostat inhibit only class I or class I and II HDAC, respectively, while many of the remaining compounds are pan-HDAC inhibitors(36–39). Another general trend we observe is that saturation of the hydroxamic acid (i.e., vorinostat) seems to result in a reduction of potency.

Of the HDAC inhibitor compounds tested, quisinostat and nanatinostat are of particular interest. Both have excellent potency (EC50s 4.5 and 23.8 nM respectively). While many compounds of the series contain an a,b-unsaturated hydroxamic acid, a potential Michael acceptor, concerning for indiscriminate reactivity, aryl hydroxamic acids quisinostat and nanatinostat both lack this feature which may mitigate some toxic potential. It is worth noting that both compounds are orally bioavailable and have been the subject of phase I and II trials in humans; thus data on pharmacodynamic, pharmacokinetic, and toxic characteristics in humans is available.

Quisinostat is a second generation hydroxamic acid-based pan-HDAC inhibitor (active in HDAC class I, II, IV)(36), however it is reported to be selective for HDAC class I when given in low doses(40). Its in vitro IC90 for *B. duncani* is 10 nM which is equivalent to a free plasma concentration of approximately 4 ng/mL. Clinical data indicates that a peak plasma concentration (Cmax) of 1.9 ng/mL and area under the curve (AUC) of 18 ng*H/mL can be achieved with a dosing strategy of 12 mg given orally (PO) thrice weekly, and with minimal side effects(40). While the Cmax appears unimpressive, it is quite possible that even with such low doses the AUC is adequate for clearance of parasitemia. Given that this dosing is a conservative approach toward mitigating toxicity for prolonged use (i.e. as chemotherapy for cancer), one could, moreover, dose this drug much more aggressively in the context of acute babesiosis treatment which would be expected to last only 7-10 days.

Nanatinostat is thought to be selective for HDAC class I-surprising given that its potency is comparable to the pan-HDAC inhibitors. Its IC90 against *B. duncani* was found to be 64 nM, equivalent to a free plasma concentration of approximately 25 ng/mL while 3x EC50 would be approximately 28 ng/mL. Clinical data indicate that a Cmax of 186 ng/mL can be achieved with a single 20 mg oral dose(41). With a single 80 mg oral dose, the protocol defined maximum tolerated dose (28 day, once-daily dosing study in humans), plasma concentrations remain above the *B. duncani* IC90 for nearly 12 hours (42). It remains to be seen if Babesia inhibition is dependent on time above MIC, AUC, or maximum serum drug concentration or if multiple doses per day are tolerable in humans.

Since growth of *Babesia* within erythrocytes is likely independent of host HDACs, quisinostat and nanatinostat likely work by inhibiting one or more *B. duncani* HDAC or, less likely, through a completely off-target mechanism. Based on BLAST search using human HDAC 8 (the most established target of quisinostat), four possible HDACs are encoded in the *B. duncani* genome. The most closely related *B. duncani* HDAC (BdWA1_002946) is conserved in the main human pathogens *B. microti* and *B. divergens* with 88% and 94% amino acid identity (96% and 97% similarity) in the known quisinostat binding pocket (43). On the other hand, sequence identity and similarity to human HDAC8 are 66% and 81%, respectively. This could partially explain the exceptionally high therapeutic index observed with these compounds and, in the case that toxicity limits the direct repurposing of quisinostat or nanatinostat to treat Babesiosis, could offer opportunities to for target-based optimization.

### Limitations and Future Directions

This work is illustrative of the value provided by an improved high throughput drug screening assay for *Babesia duncani*. The logical next step is to use it much more widely for screening of other compound libraries. A truly unbiased screening of compound libraries (i.e. those with a population predominantly noninhibitory to the parasite) would allow for calculation of a more accurate Z score. This could not be done from screening of the Bug Box because it was a collection of compounds with known antimicrobial activity, hence with a rather high pretest probability of inhibitory activity. We are currently screening several compound series known to have activity in Apicomplexa and larger, unbiased compound libraries. This should allow both for discovery of new hit compounds and for a new Z score calculation

This screen of the Bug Box represents a proof of concept for the effectiveness of our new drug screening assay. To move from assay hits to true lead compounds we would need to perform target validation and also assess drug effectiveness and safety in an animal model of disease. A murine model of *Babesia duncani* is already described(17), thus further evaluation of hits, including the HDAC inhibitor series should be a relatively straightforward enterprise. While it is a good thought experiment to consider whether the *in vitro* data and human PK data suggest that quisinostat and nanatinostat have potential for human anti-Babesia drugs, a study in mice will clearly be a necessary bridging step in order to define PK/PD relative to the MTD in humans.

## METHODS

### Culture Maintenance and Culture Preparation for Compound Screening

Frozen stocks of *B. duncani* were provided by Ben Mamoun lab at Yale University. Culture maintenance was undertaken as described by Singh et al^18^ with the following adaptations; Media, completed DMEM/F12 (cDMEM; 20% heat-inactivated fetal bovine serum (Sigma-Aldrich catalogue #12306C-500ML), 1x HT media supplement (Sigma-Aldrich catalogue # H0137-10VL), 1x L-glutamine (Gibco, catalogue # 25030-081), 1x antibiotic-antimycotic (Gibco catalogue #15240-062), and 1x gentamicin (Gibco catalogue #15710-072)), was changed every other day and cultures were kept in a modular incubator gassed with 2% O_2_, 5% CO_2_, 93% N_2_ at 37⁰C. Passage occurred once a week into fresh blood at 1-2% parasitemia, 5% hematocrit and fresh A+ blood was obtained once a week from the University of Vermont Medical Center Blood Bank. Four days before the start of drug assay testing, a 10mL, 5% hematocrit culture was infected to 2% parasitemia with *Babesia duncani*. One day before the start, media was changed, and parasitemia assessed by light microscopy examination of Giemsa-stained thin blood smears with a minimum of 500 red blood cells counted.

### Experimental Compounds

The following compounds were purchased from the respective supplier: Atovaquone (Thermo Scientific, catalogue #468762500); Azithromycin (Tokyo Chemical Industry (TCI), catalogue #A2076); Clindamycin (TCI, catalogue #C2257); Quinine (TCI, catalogue # Q0028); WR99210 (Sigma, catalogue #SML2976). The Bug Box compounds were provided by Structural Genomics Consortium/University of Toronto. Hit compounds and structurally similar series were then purchased from MedChemExpress for dose response assays (catalogue #HY-100965, #HY-10224, #HY-B0510, #HY-12163, #HY-10221, #HY-14842, #HY-13322, #HY-12164, #HY-13432, #HY-15433).

### Diluting Parasitemia Experiments

Fresh A+ blood was washed three times and a stock 10% hematocrit (HCT) solution of uninfected red cells in completed F12/DMEM media (cDMEM) was created. In a 96-well plate, pellet from a vigorously growing (≥20% parasitemia) culture was used to create a 10% hematocrit starting well that was then serially diluted five times with the stock 10% hematocrit solution. 25µL of each serial dilution was transferred to a white, clear bottom 384-well plate (Falcon^®^, catalogue #353963) in triplicates before 25µL of cDMEM was added to each well followed by 25µL of 0.6% Triton-X, 22.5µM Propidium Iodide (PI) in Phosphate Buffer Solution. The 384 well plate was then shaken for 30 seconds, incubated at 37⁰C for one hour (in the dark) then moved to room temperature in the dark. After 24-hours, the plate was read using a Synergy H1 (BioTek, Winooski, VT) with excitation = 535nm, emission = 615nm, sensitivity = 150, and readings taken from the bottom of the wells. Blood smears were made from select undiluted starting wells and stained with Giemsa, parasitemia calculated manually from a minimum of 1000 counted red blood cells. The experiment was repeated twice.

### Growth Kinetics Experiments

Experiments were performed to compare growth of *B. duncani* in 96-well and 384-well plates. For 96-well plates, a nine-point serial dilution of parasitemia was made as described above and infected erythrocytes were then seeded into a 96 well plate and incubated for 48 hours. Final hematocrit for the experiment was 5%. Blood smears were made for each point of serial dilution at both time zero and at the 48-hour mark. Smears were stained with giemsa and parasites counted, parasitemia calculated with a minimum of 1000 cells counted per slide. For 384-well plates, *B. duncani* infected red blood cells were seeded into 384 well plates at a starting parasitemia of 7.5%. At time points of 24 hours, 30 hours, 48 hours, and 72 hours parasitemia was monitored with both blood smears/manual parasite counting and PI staining and plate reading. Separate plates were used for each of the five time points. PI staining and fluorimetry of time zero point included 22 wells, the 24-hour time point included 28 wells, and all others were 27 well technical replicates.

### Z’ Score Experiments and Calculation

To calculate the Z’ score, half of a 384-well plate (154 wells) was plated with *B. duncani* infected RBCs (5% parasitemia) and treated with WR99210 (dissolved in DMSO, diluted with cDMEM) to a final concentration of 25 µM drug, 5% HCT. The other half plate was likewise infected and treated with 0.5% DMSO (final 0.25% DMSO) in cDMEM. The plate was incubated for 48-hours, stained with propidium iodide, and read as described above.

Z’ scores for the assay were calculated using the described equation in Zhang et al., 1999^15^ seen below.

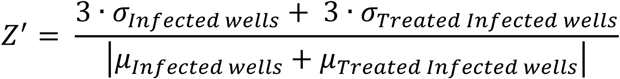

### Compound Testing

Prior to the removal of the culture(s) from the modular incubator, fresh A+ blood was washed three times with PSG+ and compounds (in DMSO stocks) to be tested were prepared in a 96-well plate by serial dilution in cDMEM. Once prepared, the culture(s) were removed from modular incubator, collected, and washed once with 7mL of cDMEM. A 10% hematocrit, 5% parasitemia stock in cDMEM was made and 25µL aliquoted into a 384-well plate with constant agitation followed by 25µL of compound dilution (quadruplicates). The positive growth control was given 25µL of 0.5% dimethyl sulfoxide (DMSO) in cDMEM (final 0.25% DMSO) and the negative growth control 25µL of 50 µM WR99210 (0.5% DMSO) (final 25 µM WR99210, 0.25% DMSO). Plates were incubated as previously described, smears of the positive growth control were made, and the plates were read as previously described. Smears were stained by Giemsa and a minimum of 1,000 RBCs were counted to determine parasitemia. Dose response assays were repeated twice.

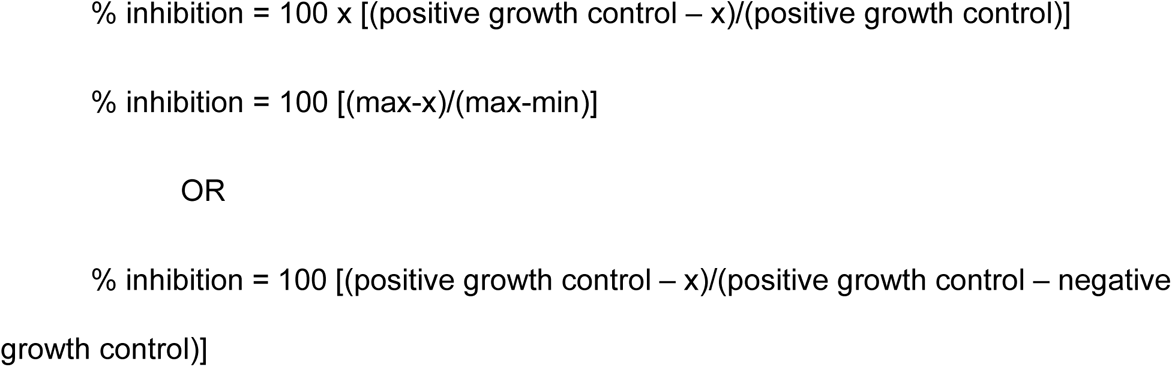

Pilot dose response experiments were performed using WR99210 (a DHFR inhibitor with known effect on Babesia duncani), DMSO vehicle, and standard of care drugs atovaquone, clindamycin, quinine, and azithromycin(20). During initial experiments positive growth control wells (infected red blood cells) were treated with 0.25% DMSO in cDMEM with a background subtraction factor established using averaged wells containing uninfected red blood cells and % inhibition calculated using formula:

We independently established the potency of WR99210 (∼8.3 nM) and then adopted the practice of using this compound as a negative growth control (minimum) in place of uninfected red blood cells. The positive growth control was treated with 25µL of 0.5% dimethyl sulfoxide (DMSO) in cDMEM (final 0.25% DMSO) and the negative growth control with 25µL of 50 µM WR99210 (0.5% DMSO) (final 25 µM WR99210, 0.25% DMSO). The plate was incubated 48 hours and the plate was stained and read as previously described. RFUs from positive and negative growth control wells were averaged (n=27). % inhibition calculation was changed to:

### “Bug Box” Compound series screen and validation

The Bug Box is a collection of 40 compounds with known antimicrobial and/or bioactivity which was provided by the Structural Genomics Consortium(SGC)-Toronto. Full list of compounds is available in supplement. They were obtained in 10 mM aliquots and screened in 384 well plates at 1 µM and 5 μM dilutions with 4 technical replicates per dilution. Hits were defined as those compounds with at least 80% inhibition. Hits were validated by dose response curves using commercially obtained compound with verified purity. The only exception to this was in the verification of compound M7 which is a proprietary compound which could be obtained only from SGC-Toronto. Dose response curves were created from two independent experiments, each with nine dilutions of drug. Panobinostat and its putative target, the histone deacetylase complex, were selected for further study with dose response curves generated for nine structurally similar compounds in effort to evaluate structure-activity relationships (SARs)

### Cyotoxicity Assays

Human HCT8 cells were grown in tissue culture flasks until a confluent monolayer was achieved. Cells were then transferred to 384 well plates where they were again grown into a confluent monolayer (approximately 48 hours of incubation). 9-point dilutions of lead compounds were prepared in 96 well plates (highest concentration 200 µM for final concentration 100 µM) and then added in 1:1 ratio to 384 well plates containing HCT8 cells. Cells were incubated with drug for 48 hours. Promega CellTiter Aqueous One Solution Cell Proliferation Assay (MTS)(PRG3580) was commercially obtained and used to ascertain cell viability with luminescence measured using a Biotek microplate reader. Cells incubated in 0.5% DMSO served as a positive growth control and comparator arm.

### Data Handling and Analysis

The raw relative fluorescent units (RFU) output from Gen5 version 1.11 were imported into Microsoft Excel for data organization and analysis. GraphPad Prism version 10.0.2 (GraphPad Software, San Diego, CA) was used to generate dose-response curves (see below) and graphs.

To generate the simple linear regression, Prism default settings were used along with the option to consider each replicate Y value as an individual point.

To generate dose-response curves, % inhibition from each experimental well calculated as described above and then % inhibition from the four replicate wells (for each compound dilution) were averaged. Each experiment was repeated at least twice with EC50 values derived from the combination of the two data sets generated. EC_50_ values were calculated in GraphPad Prism as per the following equation, with the top constrained to 100: Y = Bottom + (Top − Bottom)/(1 + 10^LogEC50^ − X × Hill slope), where Y is the percent inhibition, X is the drug concentration, and Hill slope is the largest absolute value of the slope of the curve.

## ACKNOWLEDGMENTS

Dr. Choukri Ben Mamoun and Dr. Pallavi Singh at Yale University

University of Vermont Medical Center Blood Bank

UVM LCOM Internal Grant Program

UVM TGIR-COBRE Pilot award grant

Structural Genomics Consortium-Toronto

## DATA AVAILABILITY

By publishing in the journal, the authors agree that, subject to requirements or limitations imposed by local and/or U.S. Government laws and regulations, any materials and data that are reasonably requested by others are available from a publicly accessible collection or will be made available in a timely fashion, at reasonable cost, and in limited quantities to members of the scientific community for noncommercial purposes.

## SUPPLEMENTAL FIGURES

**FIG 1s.**
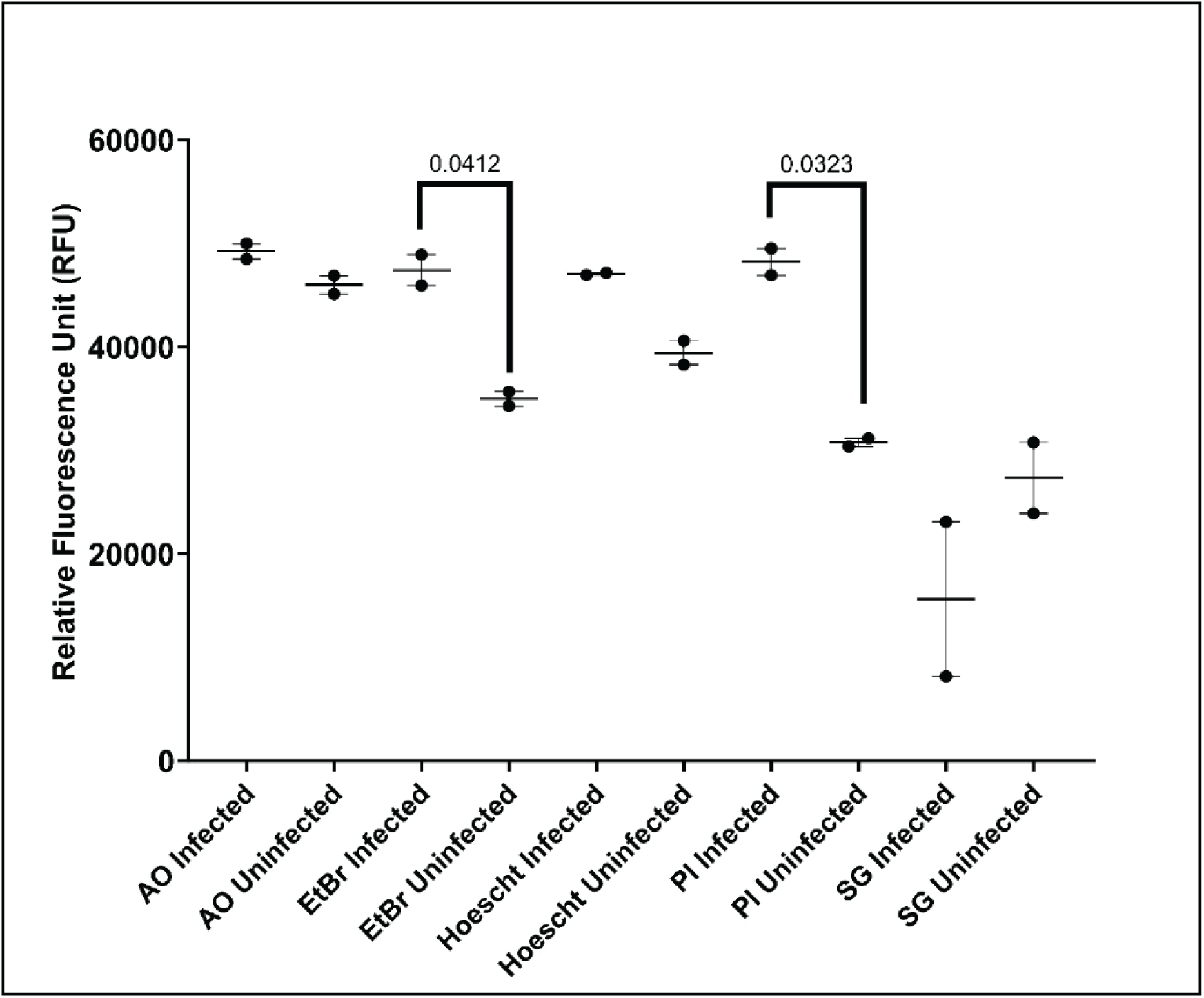
Comparison of Fluorescent Stains for distinguishing Babesia duncani-infected from uninfected red blood cells. Experiment run in 384 well plates with each point representing a single well. Abbreviations and excitations/emissions: AO = Acridine Orange (500/526); EtBr = Ethidium Bromide (301/603); Hoescht = Hoescht 33528 (352/458); PI = Propidium Iodide (535/617); SG = SYBR Green (485/530). Stains all at 1:1000 dilution. Relative fluorescence comparedbetween uninfected and infected cells using Welch’s T tests with p-values given for EtBr and PI whichwere the only stains that yielded a statistically significant result. Of the two PI had a superior signal to noise ratio.

**Fig 2s.**
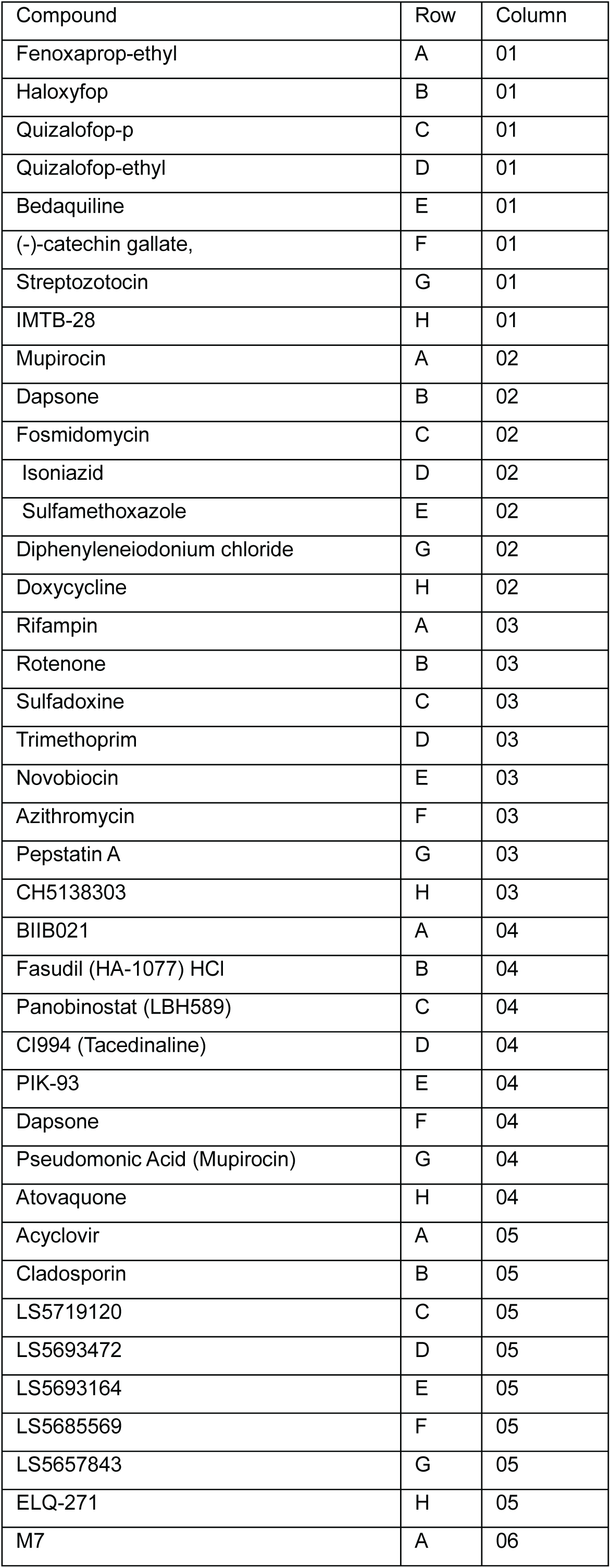
Complete list of screened antimicrobial compounds as well as location on plate. This corresponds with the coordinates given on the heatmap from FIG 3A.

**FIG 3s.**
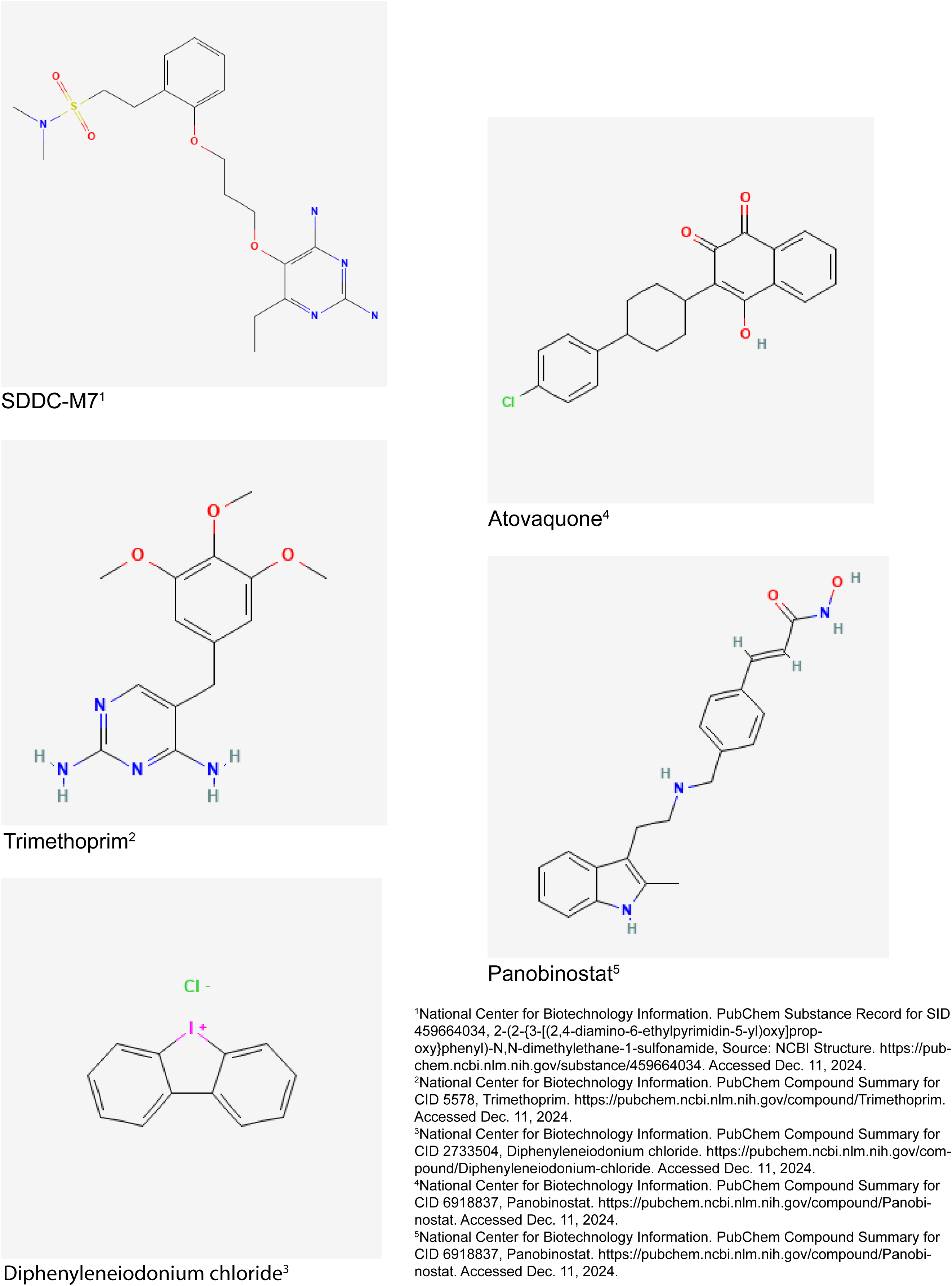
Chemical Structures of Hits from Screening of the SGC antimicrobial compound library (”Bug Box”). Structures obtained from National Center for Biotechnology Information PubChem.

